# Structural and mechanistic diversity in p53-mediated regulation of organismal longevity across taxonomical orders

**DOI:** 10.1101/2024.08.05.606567

**Authors:** Romani Osbourne, Kelly M. Thayer

## Abstract

The accumulation of senescent cells induces several aging phenotypes, and the p53 tumor suppressor protein regulates one of the two known cellular senescence pathways. p53’s regulation of senescence is however not clear. For example, p53 deficiency in some mice has been shown to rescue premature aging while others display significant aging phenotype when p53-deficient. This study seeks to elucidate, structurally and mechanistically, p53’s roles in longevity. Through a relative evolutionary scoring (RES) algorithm, we quantify the level of evolutionary change in the residues of p53 across organisms of varying average lifespans in six taxonomic orders. Secondly, we used PEPPI to assess the likelihood of interaction between p53–or p53-linked proteins–and known senescence-regulating proteins across organisms in the orders Primates and Perciformes. Our RES algorithm found variations in the alignments within and across orders, suggesting that mechanisms of p53-mediated regulation of longevity may vary. PEPPI results suggest that longer-lived species may have evolved to regulate induction and inhibition of cellular senescence better than their shorter-lived counterparts. With experimental verification, these predictions could help elucidate the mechanisms of p53-mediated cellular senescence, ultimately clarifying our understanding of p53’s connection to aging in a multiple-species context.

**Author summary:** The p53 tumor suppressor protein protects our genome from cancers by repairing DNA damage, regulating cell death and/or pushing cells to a state where they become permanently unable to divide (known as cellular senescence). An accumulation of senescent cells produces various molecular features of aging in both mouse and human cellular models–thus linking p53 to the aging process. However, the molecular mechanism by which p53 regulates aging and its structural implications on this regulation are not clear. In this study, we assessed quantitatively the evolutionary differences in p53 sequences of organisms across several taxonomical orders to determine if there is a relationship between average lifespan and sequence evolution. In addition, we used a protein-protein interaction tool to assess the likelihood of interaction between p53, or p53-associated protein, and various senescence-associated proteins across organisms of various lifespans in two taxonomic orders: Primates and Perciformes. An elucidation of p53 structural difference and mechanistic proteomic network linked to p53 regulation of cellular senescence could advance therapeutics targeting abnormal aging.

## Introduction

Usually, aging is a gradual process characterized (in human and mouse models) by molecular biomarkers such as a decrease in leukocyte telomere length, decreased levels of insulin/insulin-like growth factor (IGF-1), and increased inflammation. Several molecular mechanisms have been purported to regulate aging and determine lifespan–many of which have been linked to p53 tumor suppressor activities. In low or high-stress conditions, p53 binds to several target genes–including those that encode promyelocytic leukemia protein (PML), plasminogen activator inhibitor-1 (PAI-1), and deleted in esophageal cancer 1 (DEC1)–which then induce cellular senescence [1,2]. The link between aging and its ability to push cells to senescence is of particular interest to this study. In a senescent state, a damaged cell resists apoptosis and ceases to replicate. An accumulation of these cells triggers the aging process by creating a senescence-associated secretory phenotype (SASP) which creates a chronic inflammatory microenvironment. Chaib et. al. has shown that a programmed clearance of senescent cells delays aging phenotypes [3].

While p53 consensus sequences for most of these targets have been elucidated, few studies have explored regulatory mechanisms and structural features of p53 that could be implicated in organismal aging. Residual changes in the DNA-binding domain of several orthologs of p53 in Cetaceans have been linked to longevity [(4)]. This supports findings that p53-mediated cellular senescence could be mediated directly by DNA binding. Additionally, there has been an extension exploration of the role of the mouse double, minute two (MDM2) gene in the aging process of mice. MDM2 is the most well-studied negative inhibitor of p53 tumor suppressive activity. Wu et. al. have demonstrated that disruption of the MDM2-p53 axis accelerates aging in some mice [2]. In another study in which nearly all of their p53 N-terminal was deleted, several mice displayed high resistance to cancer, but also shorter lifespans and accelerating aging phenotypes, suggesting the importance of the MDM2-p53 praxis to the aging process [5].

However, p53’s link to organismal aging may not be easily explainable just by the MDM2-p53 axis. MDM2 has only minor regulatory effects on the levels of p53 in *Heterocephalus glaber (*the naked mole rat). And, interestingly, *H. glaber* lives an average of 30 years compared to an average of 2-4 years for most rodents. This is thought to be due in part to a hyperstable p53–the source of this stability remains largely unknown [6,7]. In another case, the African elephant–*Loxodonta africana*–despite being predisposed to cancer due to prolonged UV-radiation and large body mass lives comparably long lives and displays a significantly lower frequency of cancer when compared to humans. The unique presence of 20 copies of p53 in their genome is thought to be responsible for this [8].

The relationship between p53 and organismal longevity is thus multifaceted: changes in DNA binding, changes in N-terminal, and increases in the number of isoforms are established structural and molecular features that increase lifespan. However, whether or not these structural properties and mechanistic differences are observed across a wider range of organisms–insights that could have implications for the development of therapeutics for accelerated age-related phenotypes–is not clear. Considering the gaps in mapping out the mechanistic and structural relevance of p53 to cellular senescence and a multitude of changes in p53 that appear to correlate with longevity, investigating this relationship warrants a study of p53 orthologs in a broader evolutionary context. To address this, we develop a relative evolutionary scoring (RES) algorithm to comprehensively investigate the changes in p53 structure across organisms of various taxonomic orders and observed average lifespan to assess which residues and domains of aligned p53 sequences have evolutionary differences that could statistically explain differences in observed average lifespan.

Mapping out some mechanistic pathways of p53-induced cellular senescence across a broad sample of organisms could also be valuable to understanding the role of p53 in organismal aging. Several genes are implicated in the p53/p21 cellular senescence pathway, with some known to interact with p53, or p53 upstream regulator, following protein expression. Of interest to this study are the interactions between p53 and Smad2, Smad3, Rbl2, Npm1, MDM2, and Klf4; the interaction between MDM2 and Rpl11; and the interaction between Pras40 and Atk. These interactions either induce or inhibit cellular senescence: (a) p53 and Smad proteins: p53/Smads complex formation is required for the regulation of important tumor suppressive genes and/or cellular senescence regulatory genes. The tumor suppressor *GATA4*, which induces cellular senescence ectopically, loses this ability when Smad2 is knocked down while Smad3 complexes with p53 in its activation of the cellular senescence-inducing gene PAI-1 [9,10]. (b) p53 and Rbl2: Rbl2 is major pocket protein in the p53/p21 DNA-repair-induced cellular senescence pathway [11]. (c) p53 and Npm1: Npm1 regulates the stability and transcriptional activity of p53 via direct interaction. It inhibits p53-mediated senescence [12,13]. (d) p53 and MDM2: MDM2 ubiquitinates p53 for degradation, and thus inhibits cellular senescence abilities of the p53/p21 pathways [14]. (e) p53 and Klf4: Klf4 physically interacts with p53, causing activation of the p21 promoter [15]. (f) MDM2 and Rpl11: Rpl11 is a ribosomal protein that cooperatively binds MDM2, disrupting its ubiquitinase activity and stabilizing p53; induction of p53-dependent senescence is averted upon co-depletion of Rpl11 in HeLa cells [16,17]. (g) Pras40 and Atk: Dual phosphorylation by Atk and mTORC1 of Pras40 results in negative regulation of the Rpl11-MDM2 pathway, suppressing p53-mediated senescence [18].

We use the PEPPI protein-protein interaction prediction algorithm to assess the likelihood of interactions between these protein orthologs across organisms in the order Primates and Perciformes with varying average lifespans. Altogether, this study elucidates some structural and mechanistic diversity in p53-induced cellular senescence regulation across organisms and builds on evidence that this induction may be implicated in organismal aging.

## Results

We developed a RES algorithm to score the evolutionary changes of single residues of p53 sequences. The algorithm uses a position weight matrix and global background frequencies of amino acids observed [19,20]. 15 sequences within the taxonomic orders Primates, Rodentia, Artiodactyl, Perciformes, Cetacean, and Carnivora were first aligned in the MEGA software package using the MUSCLE alignment algorithm and then scored using RES [21]. A seventh set of sequences was aligned and scored: the top three longest-lived organisms from each of the aforementioned orders. Following residue-wise scoring of each sequence in the alignments, we calculated an r^2^ between the 15 (or 18 for the cross-order alignment) scores at each position and the observed lifespans of the 15 (or 18) organisms. We used this r^2^ to predict whether a residue could be longevity-associated, termed RES-predicted longevity-associated residue (RPLAR). After the removal of any outlier(s), we used a cutoff r^2^ of 0.3 to determine RPLARS. The RPLARs describe the residue of the unaligned ortholog of the longest-lived organism in each order. Specifications regarding the scoring algorithm and determination of this r^2^ value can be found in the Material and Methods section.

### In-order alignment revealed varying RPLARs across orders

We found no RPLAR in the artiodactyl alignment, with differences in RES where r^2^ >= 0.3 explainable only by the presence of a single outlier, *Hippopotamus amphibius kiboko*, the longest-lived organism within the alignment. The Carnivora alignment contained four RPLARs: residues 66, 70, 135, and 326 (with respect to the *Ursus maritimus* ortholog). The Cetacean alignment had two RPLARS, both in the DNA-binding domain: residues 255 and 290 (with respect to the *Balaenoptera physalus* ortholog). We found four RPLARs in the Perciformes alignment, occupying regions of the N-terminal and DNA-binding domains: residues 30, 40, 136, and 278 (with respect to the *Labrus bergylta* ortholog). The Primate alignment contained five RPLARS across the N-terminal and downstream the DNA-binding domains: residues 10, 40, 51, 289, and 296 (with respect to the *Homo sapiens* ortholog). Finally, four RPLARS were found in the Rodentia alignment: residues 12, 59, 208, and 359 (with respect to the *Heterocephalus glaber* ortholog). Fig 1 displays these findings.

**Fig 1.**
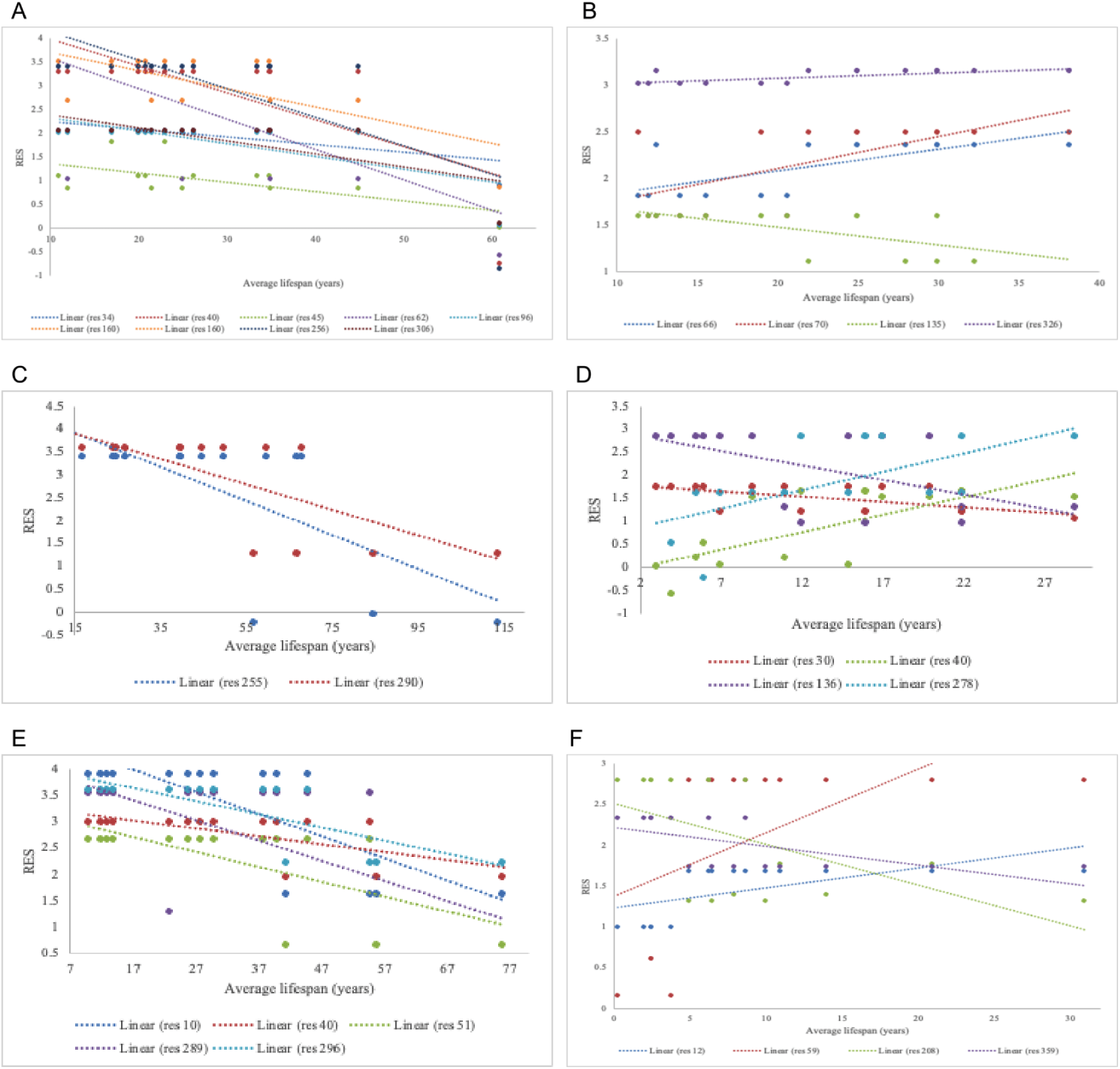
Linear plot of in-order changes in RES of RPLARS and non-RPLARs. (A) The linear difference in Artiodactyl alignment is explainable only by an outlier, thus non-RPLARs. (B) RES changes in Carnivora RPLARS: residues 66 (r^2^=0.50), 70, (r^2^=0.43), 135 (r^2^=0.37), and 326 (r^2^=0.50). (C) RES changes in Cetaceans RPLARS: residues 255 (r^2^=0.47), and 290 (r^2^=0.53). (D) RES changes in Perciformes RPLARS: residues 30 (r^2^=0.33), 40 ((r^2^=0.53), 136 (r^2^=0.32), and 278 (r^2^=0.40). (E) RES changes in Primates RPLARS: residues 10 (r^2^=0.59), 40 (r^2^=0.43), 51 (r^2^=0.43), 289 (r^2^=0.31), and 296 (r^2^=0.59). (F) RES changes in Rodentia RPLARS: residues 12 (r^2^=0.36), 59 (r^2^=0.32), 208 (r^2^=0.33), and 359 (r^2^=0.37).

### Cross-order alignment predicted most RPLARS in N– and C-terminals

Next, we conducted an alignment and RES scoring to assess the relationship between sequence diversity and observed lifespan in a cross-order context. This alignment involved the p53 orthologs of the organism with the top three highest observed average lifespans. The alignment predicted 10 RPLARs, with most of them occurring in the N– and C-terminals: residues 39, 63, 66, 69, 71, 73, 122, 305, 325, and 350 (with respect to the *Balaenoptera physalus* alignment). The greatest difference is RES for most of these RPLARs is observed between the Cetacean and Rodentia orthologs, which have the greatest difference in average lifespan (∼40 years) across the six orders. The RPLARs in this alignment also contained a greater number of amino acid substitutions that are of functionally different classes. Fig 2 depicts this.

**Fig 2.**
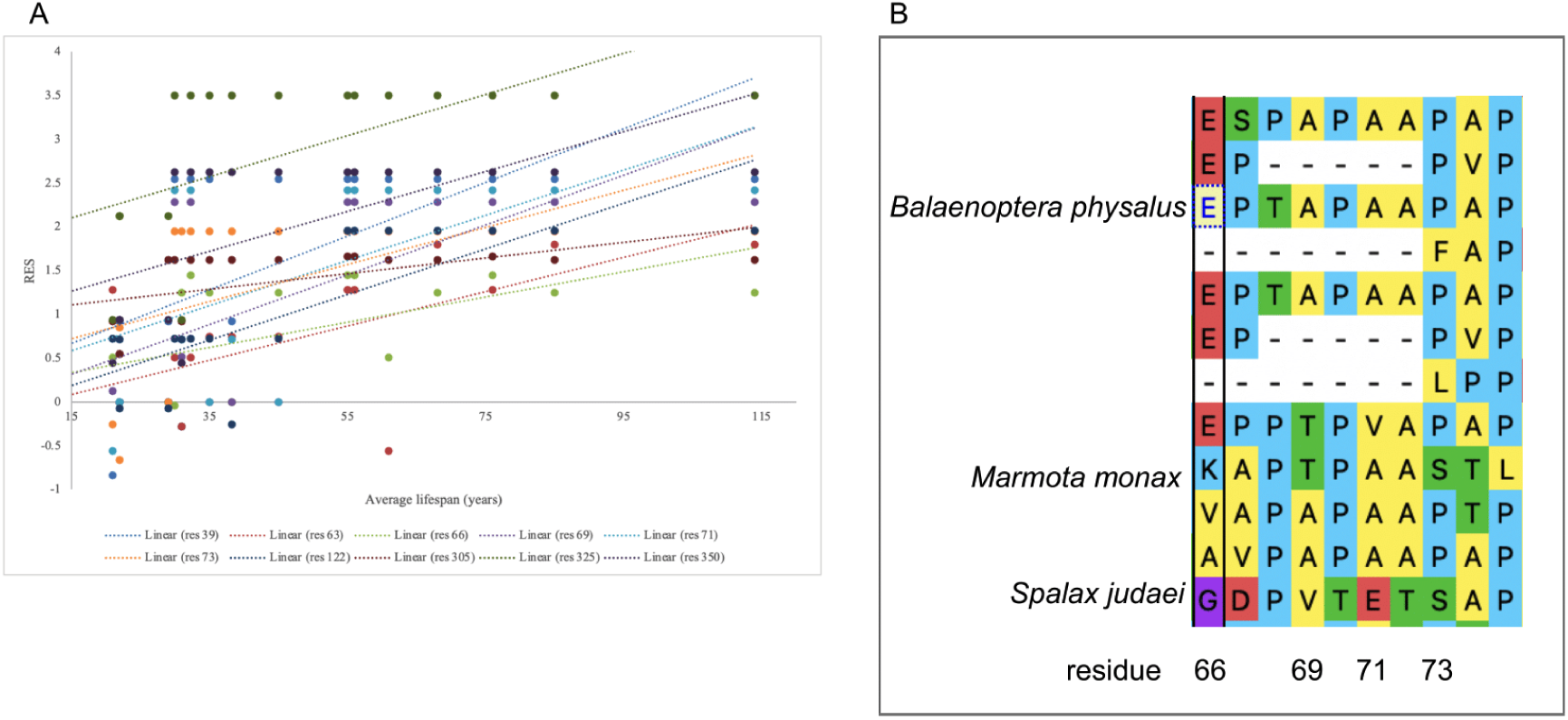
Cross-order linear plot of RPLARs and alignment. (A) RES changes in p53 orthologs of top three longest-lived organisms in each order: residue 39 (r^2^=0.42), 63 (r^2^=0.46), 66 (0.35), 69 (0.45), 71 (r^2^=0.32), 73 (r^2^=0.33), 122 (r^2^=0.64), 305 (r^2^=0.33), 325 (r^2^=0.37), 350 (r^2^=0.40). (B) Alignment of four of the ten RPLARS, showing amino acid changes between the longest-lived organism *Balaenoptera physalus* (a Cetacea) and two of the shortest-lived organisms *Marmota monax* and *Spalax judaei* (Rodents) in the alignment.

To ascertain the extent to which mutation at these identified residues (in both the in-order and cross-order alignment) affects the function of p53, we used the Sorting Intolerant from Tolerant (SIFT) algorithm. SIFT uses a sequence homology-based method to predict the likelihood of a residual substitution affecting a protein’s functionality. A SIFT score of 1 suggests that a mutation is not predicted to affect functionality while a score less than 0.05 is predicted to be deleterious; a score between these ranges is predicted to be tolerated. We compared missense mutations of p53 sequences belonging to the longest and shortest-lived organisms. If the two sequences had the same residue at a particular point, the organism with the next shortest lifespan and a residual change was tested against the longest-lived organism. The average SIFT score of the RPLARs in each order is as follows: Primates had an average score of 0.816, Carnivora had an average score of 0.715, Rodentia had a score of 0.53, Artiodactyl had an average score of 0.888, Perciformes had an average score of 0.438, and Catacean had an average score of 0.325. The average SIFT score for RPLARs from the cross-order alignment was 0.519. None of the mutations had a deleterious effect on p53 function. See Fig 3 for details of individual RPLAR SIFT scoring.

**Fig 3.**
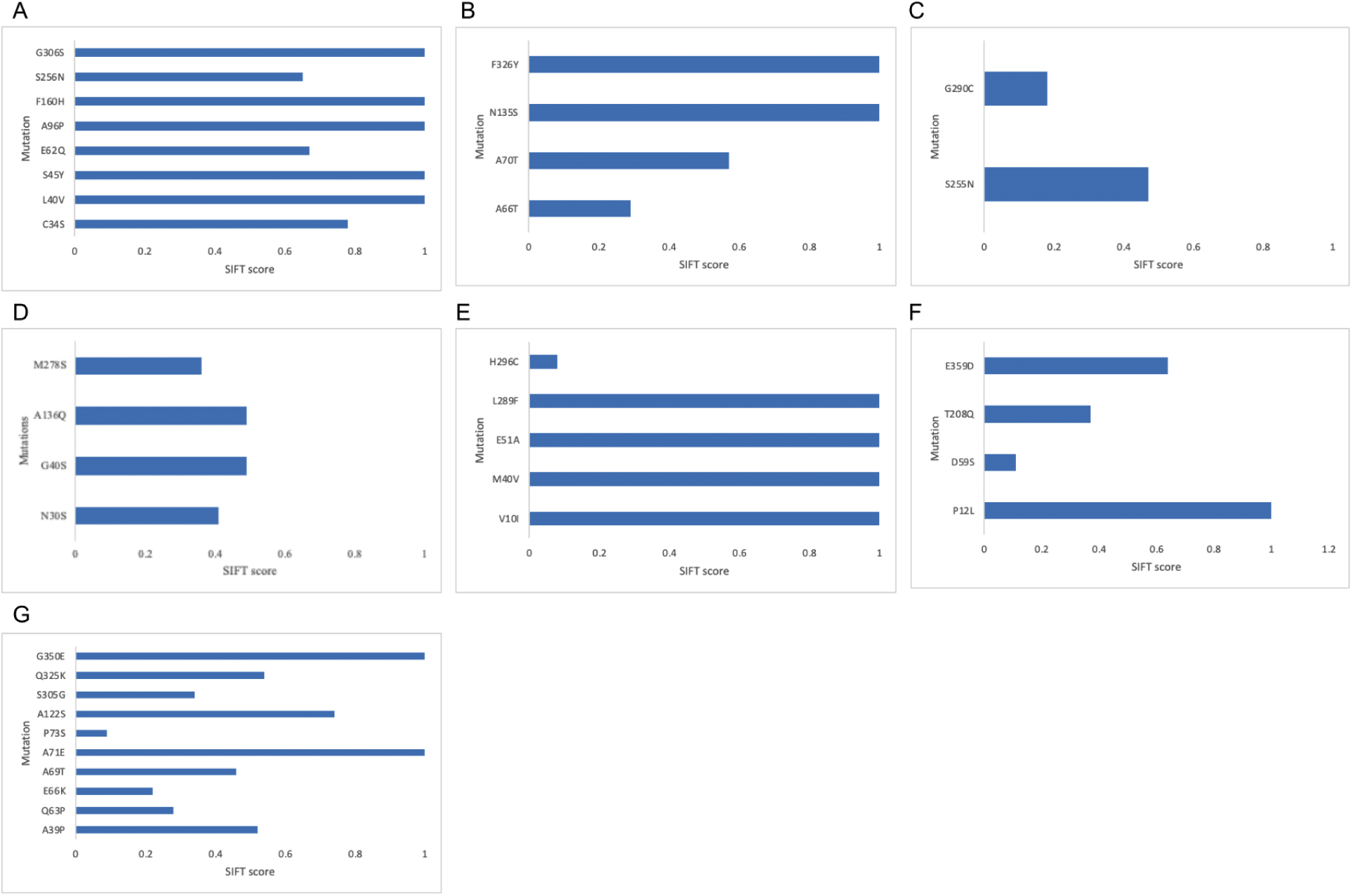
SIFT predictions of mutations at RPLARs and non-RPLARs. (A) SIFT scoring of the non-RPLARs in the Artiodactyl alignment: mutation S256N has the lowest score of 0.65. (B) SIFT scoring of the RPLARs in the Carnivora alignment: mutation A66T had the lowest SIFT score of 0.29. (C) SIFT scoring of the RPLARs in the Cetaceans alignment: mutation G290C had the lowest SIFT score of 0.18. (D) SIFT scoring of the RPLARs in the Perciformes alignment: mutation M278S had the lowest SIFT scoring of 0.36. (E) SIFT scoring of the RPLARs in the Primates alignment: mutation H296C had the lowest SIFT score of 0.08. (F) SIFT scoring of the RPLARs in the Rondentia alignment: mutation D59S had the lowest SIFT score of 0.11. (G) SIFT scoring of the RPLARs in the cross-order alignment: mutation P73S had the lowest SIFT score of 0.09.

We then used the RES and SIFT algorithms and analyses to assess the effect of potential disruption of the MDM2-praxis on the lifespans of organisms whose orthologs were used in the cross-order alignment. Using the human p53 ortholog, we assessed the r^2^ values, classification of amino acid change, and SIFT scoring of nine residues involved in the MDM2-p53 axis: residues S15, T18, S20, K370, K372, K373, K381, K382, and K386. Mutation at the tenth residue, P72, has been demonstrated to decrease lifespan in some human populations [22]. The average r^2^ (modeling the relationship between RES and average lifespan) was 0.10, with residue 72 having the highest r^2^ of 0.18. Four of the ten positions involved deletion (residues 370, 372, and 373) or no change in amino acid (residue 381), so six missense mutations were scored in SIFT. Most of the mutations proved to be deleterious, with an average SIFT score of 0.17. Mutation at residue 72 had the highest tolerance score of 0.57. In the N-terminal residues, the *Pan troglodytes* ortholog differed from that of the *Homo sapiens* while orthologs from all aligned Perciformes (*Labrus berylta*, *Perca fluvialitis*, and *Epinephelus coioides*) differed from that of *Homo sapiens* in the C-terminal residues (see Fig 4).

**Fig 4.**
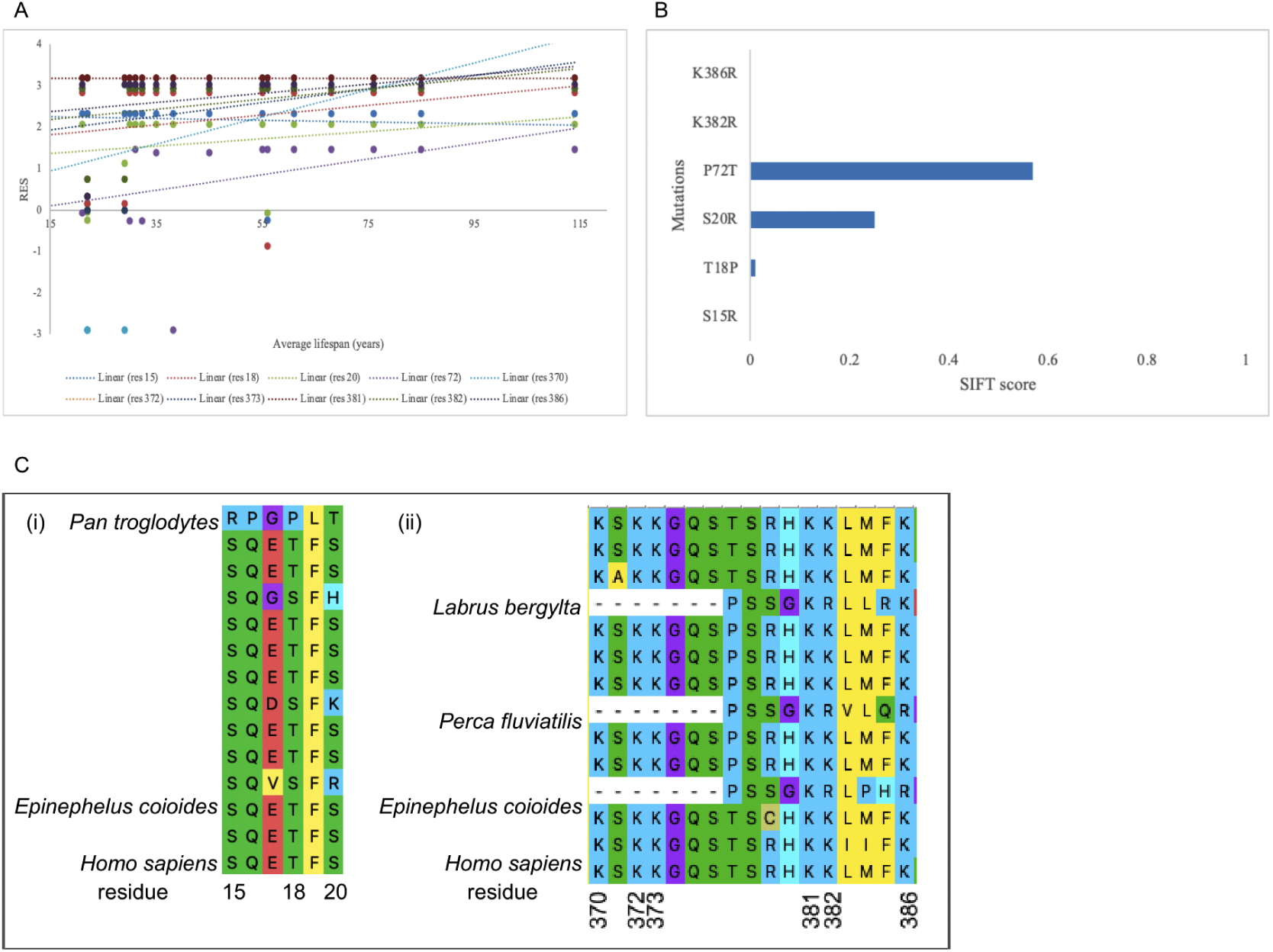
Cross-order linear plot, SIFT scoring, and alignment p53 residues involved in longevity or MDM2 regulation. (A) RES changes in p53 orthologs of top three longest-lived organisms in each order: residues 15, 18, 20, 381 (all with r^2^ < 0.1), 372, 373, 382 (all with r^2^ = 0.15), 72 (r^2^=0.18), and 386 (r^2^=0.11). (B) ASIFT scoring of the RES in the cross-order alignment: mutations S15R, K382R, and K386R all have the lowest SIFT score of 0.0. (C) Alignment of the MDM2-linked residues, showing amino acid changes at the (i) N-terminal residues primarily between the *Homo sapiens* and *Pan troglodytes* orthologs and (ii) C-terminal residues between the *Homo sapiens* and Perciformes orthologs.

### Likelihood of p53-mediated senescence interactions varied within and across orders

Several protein-protein interactions are implicated in the induction and/or inhibition of p53-mediated cellular senescence. Studies show that disruption of the p53-MDM2 induces senescence and p53-null environments may rescue aging phenotypes [2]. Considering the established role of senescence in aging, we hypothesize that the likelihood of interaction between senescence-inducing protein-protein interactions should decrease with increasing lifespan; and that the likelihood of interaction between senescence-inhibiting protein-protein interactions should increase with increasing lifespan. We used PEPPI, which combines structural and sequence information, functional association data, and a naive Bayesian classifier model to predict the likelihood of interactions (log(LR)) between eight pairs of p53 orthologs in orders Primates and Perciformes: 4 cellular senescence-inducing and 4 senescence-inhibiting interactions [23].

#### Cellular senescence-inducing interactions

For Primates, the p53-Rbl2 and MDM2-Rpl11 interactions followed our hypothesis–average lifespan increased as log(LR) decreased, though r^2^ values of 0.06 and 0.13 suggest that the relationship may not be significant. Results from all of the other senescence-inducing interactions modeled in PEPPI opposed the hypothesis and instead displayed an increase in lifespan as log(LR) increased, with the highest r^2^ value being 0.40 for interaction between p53-Klf4 in primates. Fig 5 displays these results.

**Fig 5.**
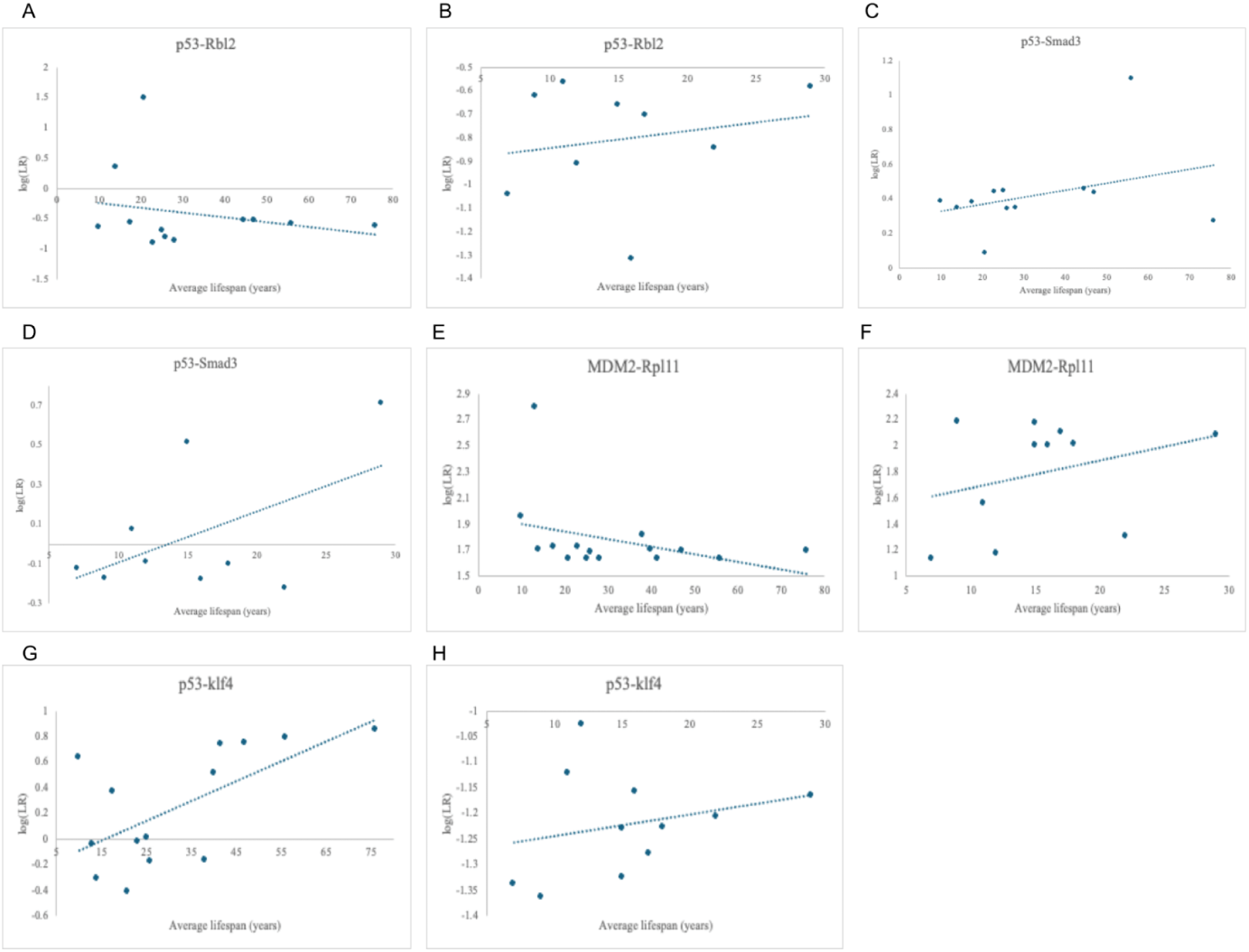
Relationship between log(LR) and average lifespan in senescence-inducing interactions. (A) Negative correlation of p53-Rbl2 in Primates (r^2^ = 0.06). (B) Positive correlation of p53-Rbl2 in Perciformes (r^2^ = 0.04). (C) Positive correlation of p53-Smad3 in Primates (r^2^ = 0.11). (D) Positive correlation of p53-Smad3 in Perciformes (r^2^ = 0.28). (E) Negative correlation of MDM2-Rpl11 in Primates (r^2^ = 0.13). (F) Positive correlation of MDM2-Rpl11 in Perciformes (r^2^ = 0.09). (G) Positive correlation of p53-Klf4 in Primates (r^2^ = 0.40). (H) Positive correlation of p53-Klf4 in Perciformes (r^2^ = 0.07).

For both p53-Klf4 and MDM2-Rpl11 interactions, we displayed the predicted interacting residues in PyMOL and found residual differences that may contribute to the difference in log(LR) observed. In the p53-Rbl2 interaction, residue H115 of p53 interacts with residue R46 of Rbl2. Alignment of the p53 sequences reveals that the ortholog of one of the shortest shortest-lived Primates, *Theropithecus gelada*, uniquely has a lysine residue at aligned position 115. In the MDM2-Rpl11 interaction of the shortest-lived organism (and that with the highest log(LR)), Rpl11 is predicted to interact with MDM2 from several residues in its N-terminal. For the organisms with the lowest log(LR) and two of the highest average lifespans, *Pan troglodytes* and *Pan paniscus*, Rpl11 is predicted to interact with MDM2 via approximately 30 residues downstream its N-terminal domain (see Fig 6).

**Fig 6.**
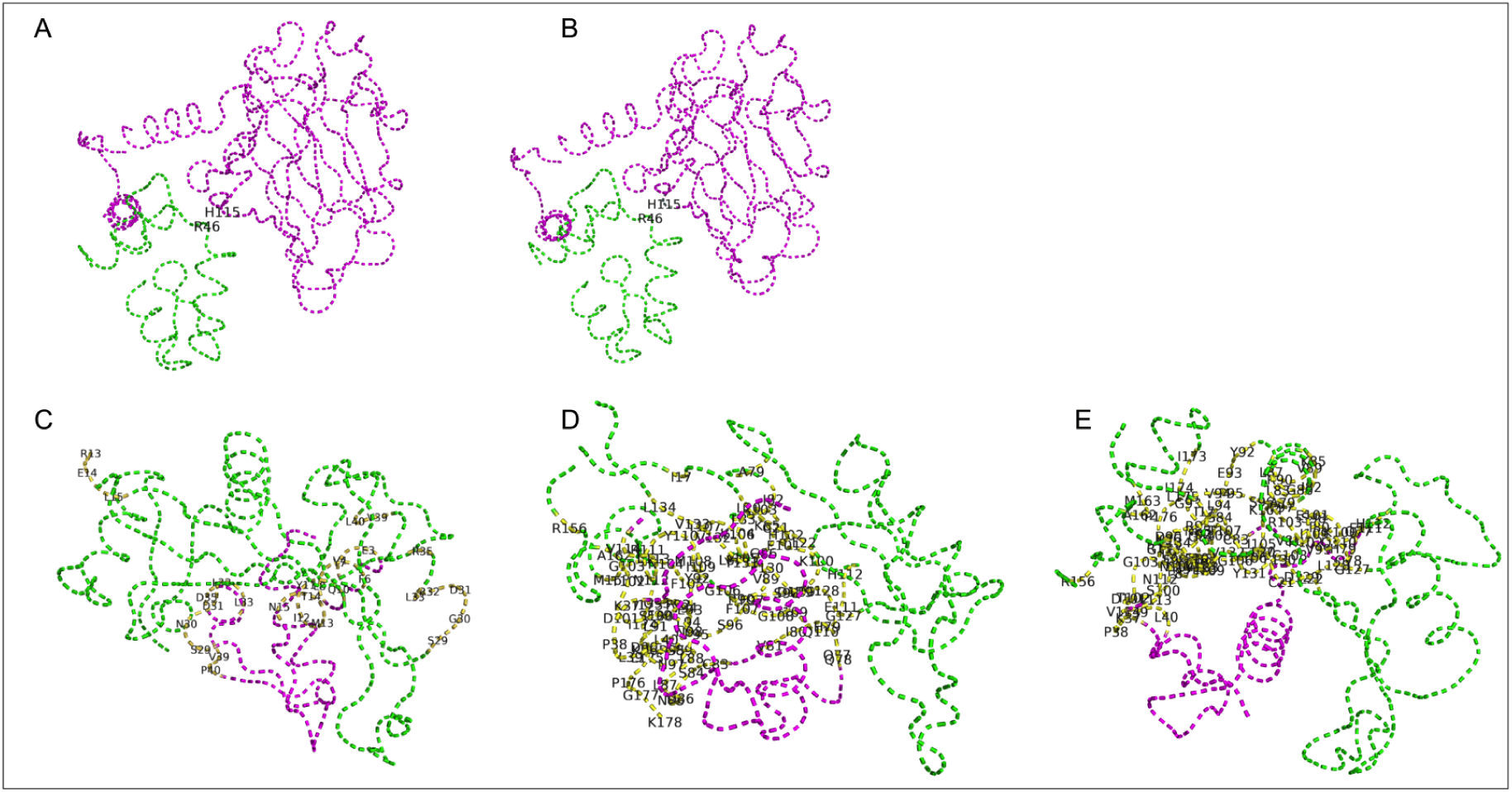
PyMOL-predicted interacting residues. (A) Model of human p53-Klf4 ortholog with interaction between p53 His115 and Klf4 Arg46. (B) Model of *Pan troglodytes* (Chimpanzee) p53-Klf4 ortholog with interaction between p53 His115 and Klf4 Arg46. (C) Model of *Carlito Syrichta* (Philippine tarsier) MDM2-Rpl11 ortholog with Rpl11 interacting from its N-terminal residues. (D) Model of *Pan troglodytes* MDM2-Rpl11 ortholog with Rpl11 interacting from downstream its N-terminal residues. (D) Model of *Pan paniscus* (Bonobo) MDM2-Rpl11 ortholog with Rpl11 interacting from downstream its N-terminal residues.

#### Cellular senescence-inhibiting interactions

For both Primate and Perciformes, the p53-Smad2 and p53-MDM2 interactions conformed to our hypothesis with the greatest strengths: average lifespans increase with increasing log(LR) with r^2^ values of 0.13 and 0.12 for the p53-Smad2 interactions and r^2^ values of 0.21 and 0.13 for the p53-MDM2 interactions, respectively. The likelihood of Pras40-Akt and p53-Npm1 interactions in both the Primates and Perciformes are virtually constant or have low r^2^ values (< 0.03) across average lifespans (Fig 7). PyMOL found no detected interacting residues in any of the

**Fig 7.**
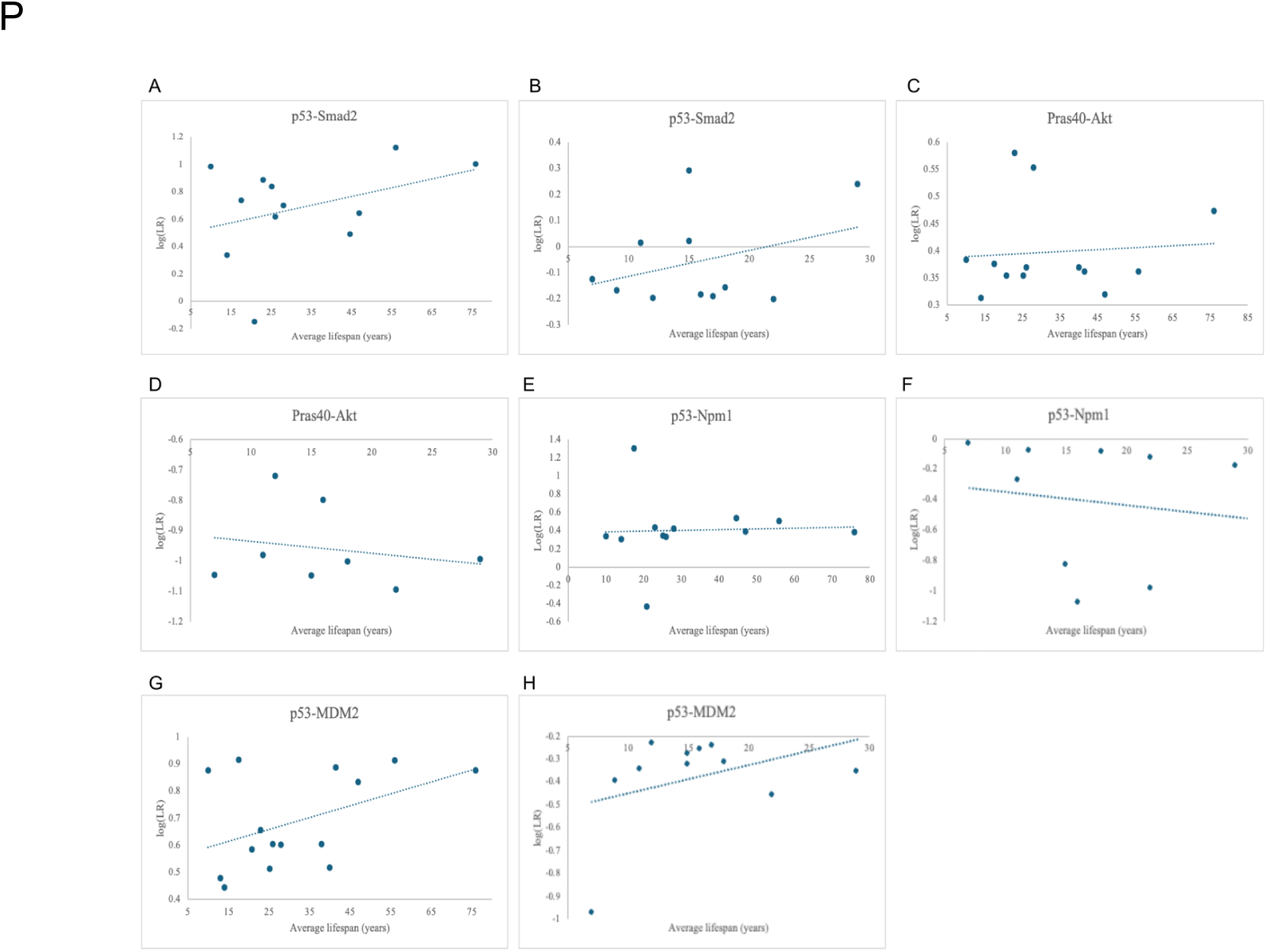
Relationship between log(LR) and average lifespan in senescence-inhibiting interactions. (A) Positive correlation of p53-Smad2 in Primates (r^2^ = 0.13).(B) Positive correlation of p53-Smad2 in Perciformes (r^2^ = 0.12). (C) Positive correlation of Pras40-Akt in Primates (r^2^ = 0.007). (D) Negative correlation of Pras40-Akt in Perciformes (r^2^ = 0.044). (E) Positive correlation of p53-Npm1 in Primates (r^2^ = 0.001). (F) Negative correlation of p53-Npm1 in Perciformes (r^2^ = 0.02). (G) Positive correlation of p53-MDM2 in Primates (r^2^ = 0.21). (H) Positive correlation of p53-MDM2 in Perciformes (r^2^ = 0.13).

For all the senescence-inhibiting interactions, the average log(LR) for Primates WAS greater than that for Perciformes–with all of the Primate interactions predicted as interacting while those in Perciformes were predicted as non-interacting. The difference in the senescence-inducing interactions between the Primates versus Perciformes differed from this observation. On average, the p53-Rbl2 interaction for the Primates orthologs was predicted as non-interacting, similar to the Perciformes ortholog with a higher non-interacting likelihood. Overall, the differences in the likelihood of senescence-inducing interactions between Primates and Perciformes are significantly less than those of senescence-inhibiting interactions (Fig 8).

**Fig 8.**
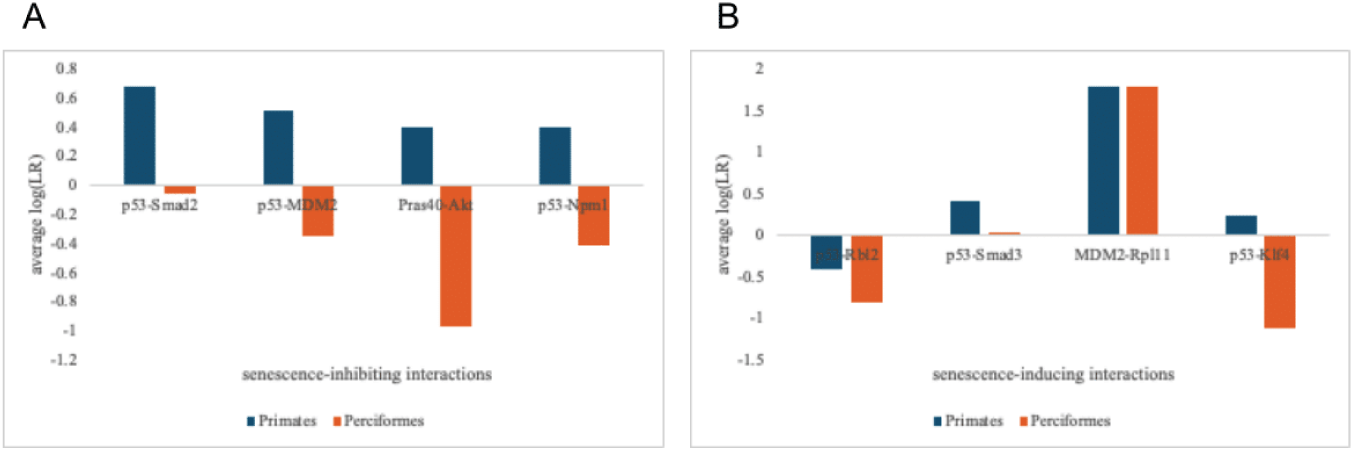
Average log(LR) for the 8 interactions. (A) Average log(LR) for senescence-inhibiting interactions in Primates (n=12-15) and Perciformes (n=8-11). (B) Average log(LR) for senescence-inducing interactions in Primates (n=12-15) and Perciformes (n=9-11).

## Discussion

Senescence regulatory activities by the p53 tumor suppressor protein have been linked to aging on the molecular level, and a few studies have implicated this link to organismal aging–mainly in mouse models. In this study, we created a relative evolutionary scoring (RES) approach that scored aligned p53 sequences in search of residues that display a relatively strong statistical relationship (r^2^ >= 0.3) to differences in observed lifespans across organisms in six taxonomical orders. In addition, we examined eight protein-protein interactions (of Primates and Perciformes orthologs) that are implicated in the p53/p21 cellular senescence pathway. We used the PEPPI protein-protein predictor to assess the likelihood of interaction of four senescence-inducing and four senescence-inhibiting interactions.

### p53 structural diversity linked to varying lifespans

Our RES findings suggest that the p53 tumor suppressor residues that influence the difference in observed lifespan vary across the six taxonomical orders. The Artiodactyla alignment contained no RPLAR; the Carnivora and Rodentia alignments both contained two RPLARs in their N-terminal, one in their DNA-binding domains, and another in their C-terminal; the Perciformes alignment contained two RPLARs in both their N-terminal and DNA-binding domain while the Primates alignment contains three RPLARs in the N-terminal and two in its DNA-binding domains. The Cetacean alignment uniquely contains two in its DNA-binding domain.

This difference in the domains in which RPLARs are found suggests that cellular senescence may be variably regulated across orders. In mouse models, the N-terminal contains the binding site for the most well-documented inhibitor of p53–the mouse double minute two (MDM2) protein. Disruption of the MDM2-p53 axis in some mice induces premature senescence and thus accelerated aging [24]. We hypothesize that p53 orthologs in Carnivora, Perciformes, Primates, and Rodentia may depend on similar inhibitory mechanisms at their N-terminal residues in regulating cellular senescence.

Differences in the p53 DNA-binding domain of Cetaceans have been linked to longevity [4]. Our results support this finding and suggest that DNA binding may also be an important regulator of longevity in all five orders with RPLARS in the DNA-binding domain. It is well known that p53 binds and upregulates several senescence-associated genes, hence, it is feasible why the DNA-binding domain may be structurally important in the regulation of longevity. Also, the N-terminal of p53 interacts with the DNA-binding domain to regulate the protein’s stability [25]. This may explain the presence of RPLARS in the N-terminal of p53 orthologs in Carnivora, Perciformes, Primates, and Rodents. We propose that the p53 ortholog in organisms in these orders may depend on interactions between their DNA-binding and N-terminal residues in their regulation of cellular senescence. SIFT scoring qualifies missense mutations of all RPLARS as tolerated, but an average scoring of 0.56 (on a scale of 0-1.0, where 1.0 predicts a change with the greatest tolerance) suggests that some mutations are causing some functional change. Whether or not these changes are affecting the regulation of longevity needs to be further explored.

Unlike the in-order alignments, from which most RPLARs are predicted to be in the DNA-binding domain, the cross-order alignment predicts most RPLARS to be in the N-and C-terminals. In a cross-order context, changes in the N-terminal may be linked to p53’s downstream regulation by factors such as post-transcriptional modifications or inhibitory oncoproteins such as MDM2. Changes in the C-terminal may also be crucial to p53’s regulation of longevity. Mouse p53 binds DNA and regulates genes as a tetramer [26]. The C-terminal is crucial for the tetramerization and stabilization of p53-DNA complexes, thus it is feasible that amino acid substitutions that significantly alter the chemical environment of C-terminal residues may affect p53’s ability to tetramerize and stably bind DNA [27].

The effects of the disruption of the MDM2-p53 axis on organismal longevity remain unclear, though some of our results point to its involvement. p53 residues S15, T18, and S20 serve as sites of phosphorylation by either ATM or Chk2. Phosphorylation of these residues causes MDM2 dissociation from p53, which enhances p53 transcriptional activities [28]. These residues are virtually constant in the cross-order alignment, with the only amino acid change observed in the *Pan troglodytes* (Chimpanzee) ortholog. While the S15R and T18P mutations (the human ortholog has S15, T18, and the chimpanzee ortholog has R15, P18) are predicted to be deleterious, the chimpanzee remains one of the longest-lived Primates. Our RES algorithm does not classify these sites as RPLARs as they both have r^2^ < 0.1. We propose two hypotheses for this observation: (1) these MDM2-linked residues in the N-terminal may have no or negligible effect on organismal aging and other sites within the N-terminal may be involved in MDM2-p53 association or/and (2) these MDM2-linked residues in the N-terminal have an effect on organismal aging and whether accelerated aging phenotypes is observed (as a result of disruption to the MDM2-p53 axis) depends on downstream mechanisms such as the genomes’ ability to clear senescent microenvironments. Residue 72 follows, partly, the first hypothesis: its RES most strongly explains the difference in average lifespans, and the P72 SNP increases the lifespan of individuals by 6% over a 12-year period compared to R72 [22]. Interestingly, the R72 SNP influences increased phosphorylation of p53 S15 and transactivation of p21 compared to P72 [29]. Thus, the P72 SNP could help maintain the MDM2-p53 axis and inhibit the p53/p21 cellular senescence pathway, and thus reduce accelerated aging. Still, MDM2 regulation of p53-mediated cellular senescence appears to conflictingly depend on S15 phosphorylation. Our N-terminal RPLARS may play a similar role in this regulation.

The RES changes in the C-terminal residues have higher r^2^ values (r^2^ > 0.1) across the aligned orthologs than those in the N-terminal, though they are not RPLARs. Residues K370, K372, K373, K381, K382, and K386 are sites for acetylation by CBP/p300 and ubiquitination by MDM2 [30]. Acetylation of these residues is thought to prevent ubiquitination by MDM2, thus enhancing p53 stability [31]. As MDM2 ubiquitination at these residues triggers p53 degradation, there is a constant play between MDM2 and CBP/p300 to degrade or stabilize p53. The Perciformes in the alignment uniquely contain either a deletion or arginine substitution at these residues, and all missense mutations are predicted to be deleterious by SIFT. Out of the six orders, and besides Rodentia, Perciformes have the lowest average lifespan. The changes in their sequence at these acetylation and ubiquitination residues appear to have a substantial effect on the MDM2-p53 axis and potentially their longevity. With potential disruption to acetylation, p53 orthologs of Perciformes may not be appropriately stabilized when needed, and could potentially expose their genome to excessive damage, predisposing them to shorter lifespans [32]. And, with potential disruption to ubiquitination, p53 levels may remain largely unregulated, triggering uncontrolled activities such as the p53/p21 pathway which could enhance the accumulation of senescent cells, and thus accelerate aging.

### Mechanistic diversity in induction and inhibition of p53-mediated cellular senescence

From the PEPPI predictions, the observed likelihood of senescence-inducing and senescence-inhibiting protein-protein interactions across primates and Perciformes reifies the diversity in p53 regulation of senescence that our RES study suggests. The established role of cellular senescence in driving aging at the molecular level and findings that mouse models with unregulated p53 display accelerated aging phenotypes, led us to the hypothesis that, on an organismal level, the relative likelihood of p53-senescence-regulating protein-protein interactions may be implicated in aging.

Our results of 8 such interactions complicate this. For the interactions that are found to induce p53-mediated cellular senescence, only the p53-Rbl2 and MDM2-Rpl11 interactions conformed, with low r^2^ values of 0.06 and 0.13 respectively, to our hypothesis that average lifespan would increase if the likelihood of interaction decreases. Similarly, for the interactions that are found to inhibit p53-mediated cellular senescence, the p53-MDM2 and p53-Smad2 interactions conformed with the hypothesis that an increase in log(LR) would be observed as average lifespans increase. While the study predicts the p53-MDM2 interaction to be the strongest inhibitor p53, the consistently low r^2^ (<0.5) between log(LR) and average lifespan suggests that the mere likelihood of induction or inhibition of a single pair of protein-protein interactions may not fully or accurately explain observed differences in average lifespans within a taxonomic order. Considering the established complexity of p53 interactions in a multitude of physiological and metabolic pathways, its role in senescence within an order may be more integrative and compounded. Additionally, results from the Primates and Perciformes p53-Smad3, p53-Klf4, and Perciformes MDM2-Rpl11 interactions (which all deviate from our initial hypothesis) suggest that cellular senescence, through these senescence-inducing these pathways, could be enhancing longevity. Cellular senescence is a requisite for a plethora of physiological processes such as wound healing and embryonic development, and hence why its induction may be necessary [33].

While the intra-order relationship between the likelihood of interaction and the average lifespan is quite subtle, this relationship when viewed from a cross-order context appears much clearer. For the senescence-inhibiting interactions, the Primate orthologs have significantly higher likelihoods of interaction than the Perciforme orthologs. We speculate that a feature of Primates that evolved longer lifespans (compared to Perciformes) is the ability to inhibit p53-mediated cellular senescence. Perciformes compete better in senescence-inducing interactions, though Primates still average better likelihoods of interactions. This, coupled with the established “good” and “bad” roles of cellular senescence suggests that organisms may need to create an appropriate balance between induction and inhibition of senescence to maintain healthy aging.

### Conclusions & Future Directions

We found variations in the p53 tumor suppressor residues in Primates, Rodentia, Artiodactyl, Carnivora, Cetaceans, and Perciformes that could be linked to organismal aging. Our RES algorithm prioritizes p53’s direct binding capabilities, but studies have demonstrated that regulation of and by p53 could be mediated allosterically [34]. The results from our PEPPI study give us new insight to begin mapping out a p53-mediated cellular senescence proteomic landscape. Several computational and experimental approaches could be employed to assess the effect(s) of this diversity in both the structure and mechanisms of p53 across organisms. Site-directed mutagenesis of p53 orthologs from organisms in each order, followed by an affinity binding assay could be conducted. If mutations are determined to cause significant functional change, then model organisms could be created with specific strains of p53, and their lifespans compared to those with the wild type. As Cetaceans and Rodents have the greatest difference in RES in the cross-order alignment, *in vitro* binding assays could be carried out to quantify binding between known N-terminal interacting proteins such as MDM2 and the p53 orthologs of rodents and cetaceans. To compare the degree of tetramerization of p53 in organisms of these orders, vapor diffusion could be employed to crystallize orthologs from these organisms. Molecular dynamics could also be used to assess potential allosteric pathways that our RPLARs may be implicated in as well as to gain insight into the thermodynamic and mechanistic changes that occur as a result of RPLAR mutations.

Tumor suppressor p53-mediated cellular senescence is variably mediated across orders. Longevity may be linked to these variations via p53’s tight balancing of induction and inhibition of cellular senescence. Further investigation of p53’s role in this balancing play is required. Altogether, our study reifies the established complexity of p53’s regulating of longevity and provides crucial molecular information that, through experimental verification, could enable the development of various models of p53 regulation of aging, ultimately making way for therapeutic targets to treat various accelerated aging-related diseases.

## Materials and Methods

### Observed Average Lifespan Data and Protein sequences

The observed average lifespan for each organism was obtained from the Animal Diversity Web (https://animaldiversity.org/; accessed on 13 January 2024), the AnimalLifeExpectancy (AnAge, UMICH, Max Planck, PanTHERIA, Arkive, UKC, AKC) databases as well as from Smithsonian National Zoo Database. Organism in each order was selected such that the order had a wide range of observed average lifespans. Protein sequences in RES, SIFT, and PEPPI studies for all organisms were obtained from the UniProt database (https://www.uniprot.org/; accessed on 15 October 2023) or the NIH’s GenBank database (https://www.ncbi.nlm.nih.gov/genbank/; accessed 12 October 2023). See Tables 1-6 in the Supporting Information section for a list of the organisms, their observed average lifespans, and UniProt or GenBank accession numbers for protein sequences used throughout the study.

### RES algorithm

Using the MUSCLE algorithm, p53 sequences were aligned by taxonomical order, and then the position weight matrix and background frequency values were calculated from each alignment. The position weight matrix is calculated by first counting the frequency of each amino acid (or gap) at every position in the alignment, then normalizing the count to the probability that a particular amino acid will be found at a particular position in the alignment. Included in this count of frequency is a pseudo-count to avoid a count of zero–an occurrence unlikely in naturally occurring protein sequences. We then calculated a corresponding dictionary of the overall (background) frequency of each amino acid (or gap) in the alignment. Including background frequency information in multiple sequence scoring algorithms improves scoring and identification of functional sites. Following this, we used our position weight matrix and background frequencies to score each position of individual aligned sequences by taking the log odds ratio of observed frequency to background frequency. Log-odds scoring has been used in most pair-wise scoring, as it formalizes differentiating between more functional and less functional positions [35]. We used the MUSCLE default gap opening and extension penalties of –2.90 and 0, respectively in our scoring. Link to RES github repository can be found in the Supporting Information section.

### Assessing the functional significance of RES scoring

With respect to the observed average lifespan, a coefficient of determination (r^2^) was calculated using Excel’s CORREL and POWER functions for each residual position. This r^2^ is used to determine the fraction of the observed lifespan that is explainable by the difference in RES scoring at that position. An r^2^ > 0.3 was hypothesized to predict a residue as longevity-associated. We used a combination of the relative r^2^ values across all residues and the frequency of amino acid changes observed at each position in estimating this r^2^ cutoff. The average r^2^ of each position across all six in-order alignments was found to be 0.03, with most positions having conserved residues across the aligned orthologs. We observed that positions with r^2^ of about 0.2 had amino acid substitutions, though the substitution was often limited to a single outlier. We therefore selected for an r^2^ of or greater than 0.3 and then checked the RES at those positions for outliers. If no outlier was found, we determined that that position (relative to longest-lived ortholog) was an RPLAR and reported that r^2^. If an outlier was found, we removed the outlier from that position’s sample and recalculated the r^2^, again using a cutoff of 0.3 to determine if a residue was an RPLAR.

To assess the functional correctness of this hypothesis, we employed SIFT [36]. We compared missense mutations of p53 sequences belonging to the longest and shortest-lived organisms. If the two sequences had the same residue at a particular point, the organism with the next shortest lifespan and a residual change was used in the prediction. Accession numbers of the p53 ortholog of the longest-lived organism in each order can be found in S1-S6 Tables in the Supporting Information section. Missense mutations in the ortholog of (one of the) shortest-lived organisms in each order is determined from MEGA alignment.

### PEPPI: predicting the likelihood of p53-induced cellular senescence protein-protein interactions

PEPPI, developed by the Zhang Lab at the University of Michigan Medical School, effectively uses structural and sequence similarity, functional association data, and the Naive Bayesian classifier model to predict the likelihood of interactions between proteins [13]. This study scored protein-protein interactions using its web server [23]. PEPPI runs were conducted on the interactions between p53 and Smad2, Smad3, Rbl2, Npm1, MDM2, and Klf4; the interaction between MDM2 and Rpl11; and the interaction between Pras40 and Atk of the Prtimates and Perciformes orthologs. Accession numbers of the sequences used can be found in S1 and S2 Tables of the Supporting Information section.

### PyMOL: visualizing protein-protein interaction predictions

PEPPI outputs a PDB model of predicted protein-protein interaction (or non-interaction). We used the InterfaceResidues program in PyMOL with default settings and arguments to display the predicted interacting residues of the model [37].

## Acknowledgments

We would like to thank the Molecules to Medicine Consortium members: special thanks to Professors Micheal Weir, David Beveridge, and Mathew Young for their continuous feedback throughout this project. We would also like to extend thanks to In Sub Han of the Thayer Lab who aided in member onboarding and mentorship in the lab. Thanks also to Henk Meij who maintains Wesleyan’s High Performance Computing cluster on which part of this project was conducted. The NIH grant R15 GM128102-02 funded this study.

## Supporting Information

**S1 Table.**
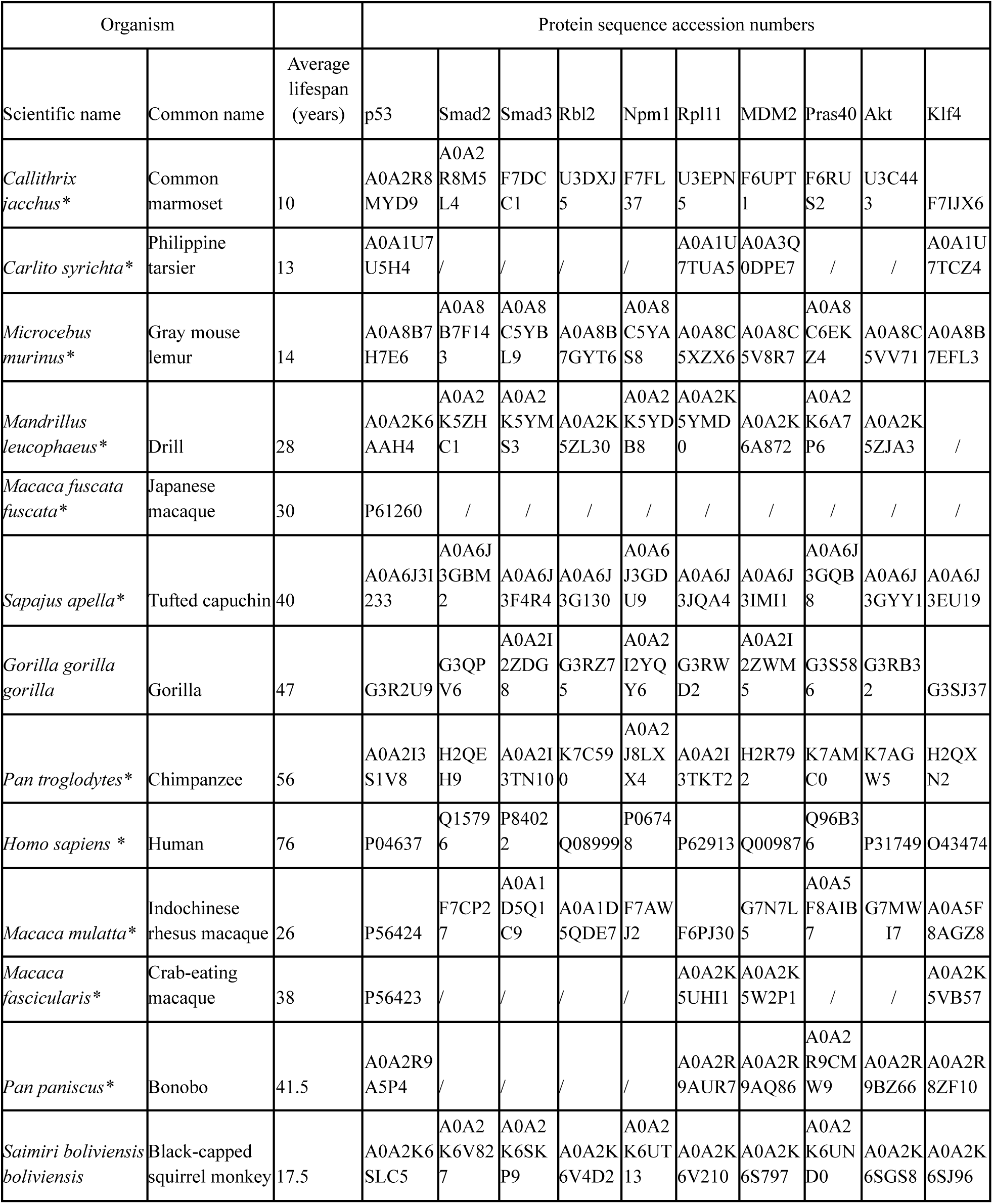
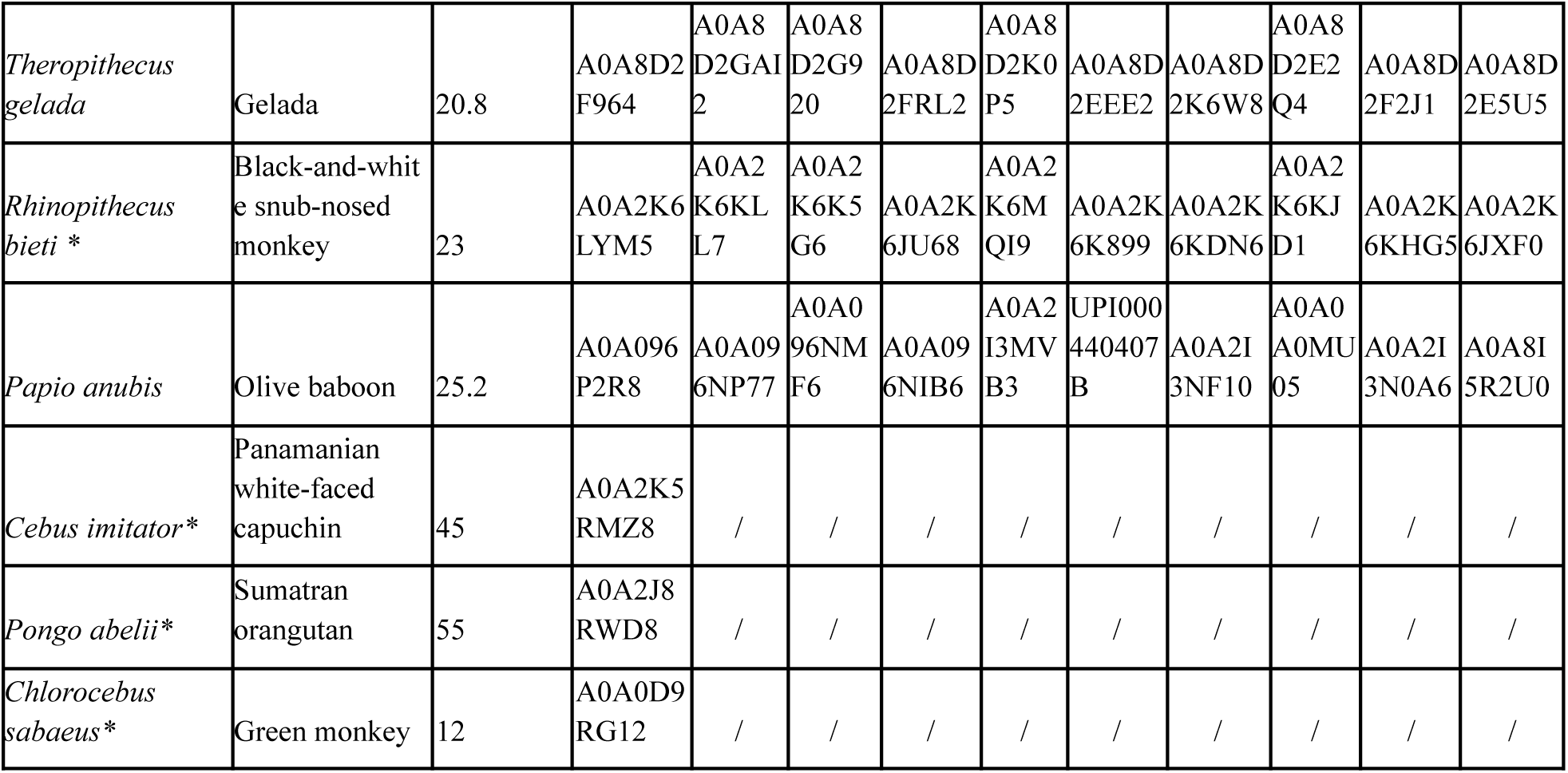
Primates included in the study. Organisms in order Primates included in the RES, SIFT, and/or PEPPI studies, their average lifespans, and UniPro or NCBI GenBank’s protein accession numbers. * indicates those organisms whose p53 sequences are used in alignment for RES study.

**S2 Table.**
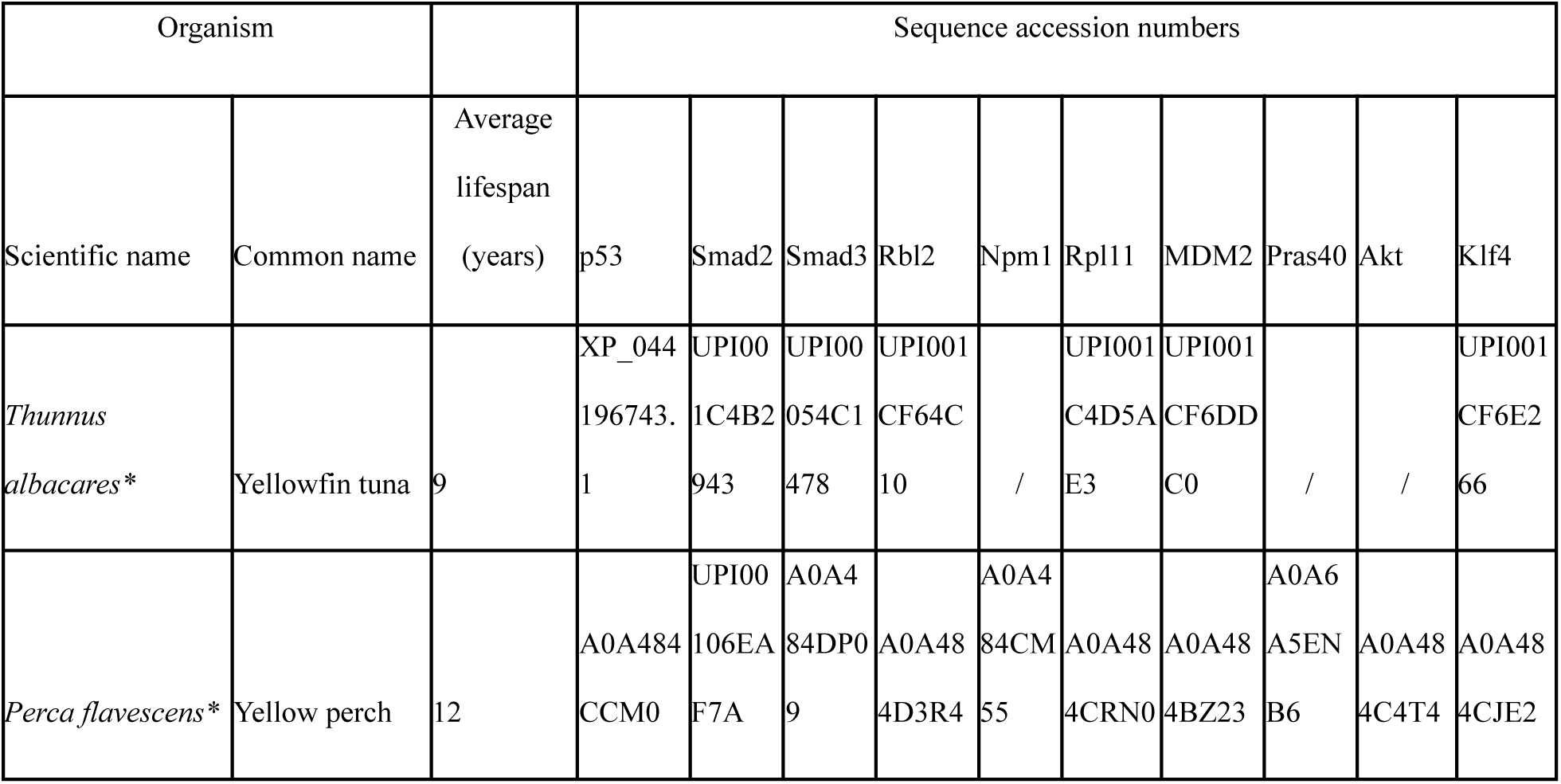
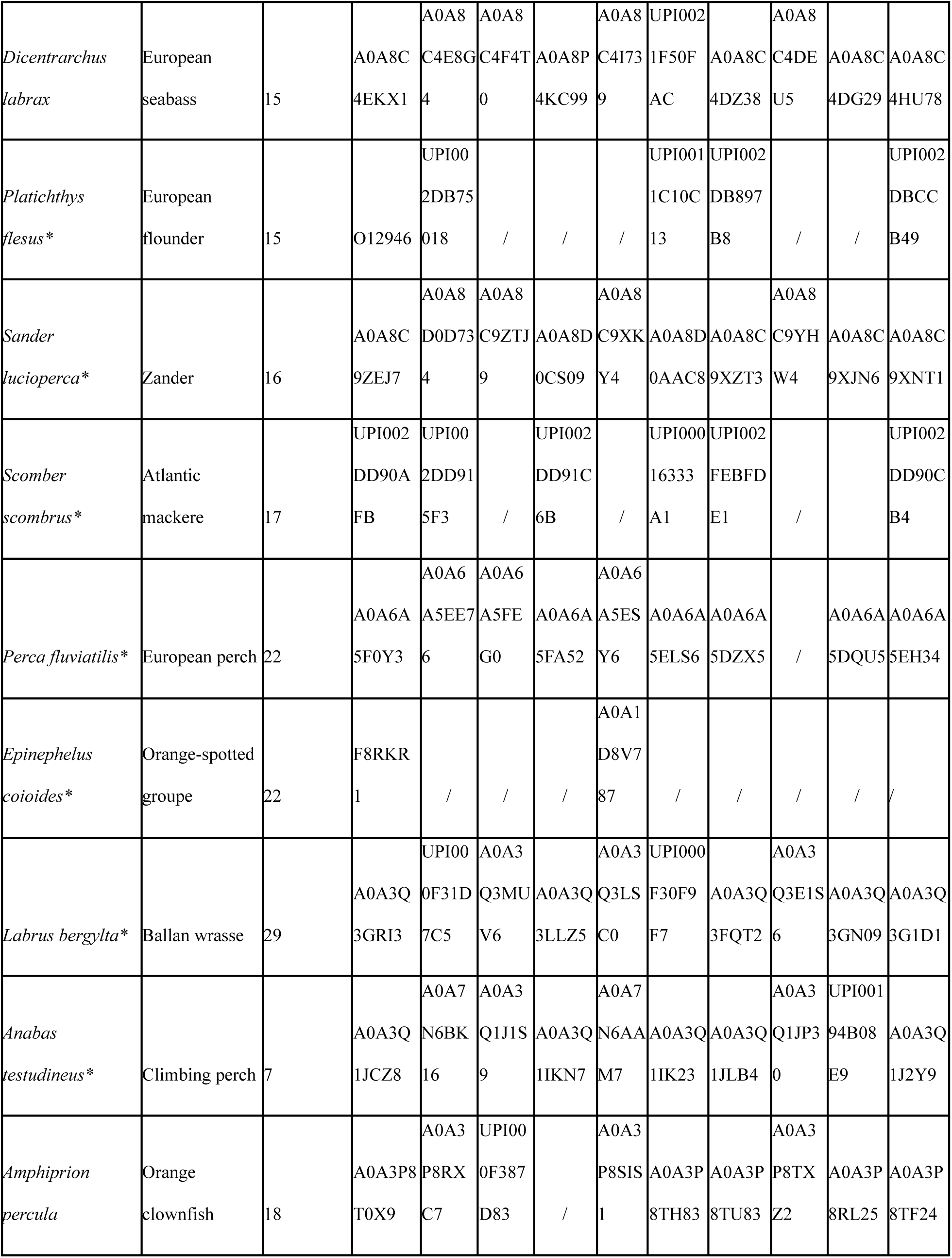
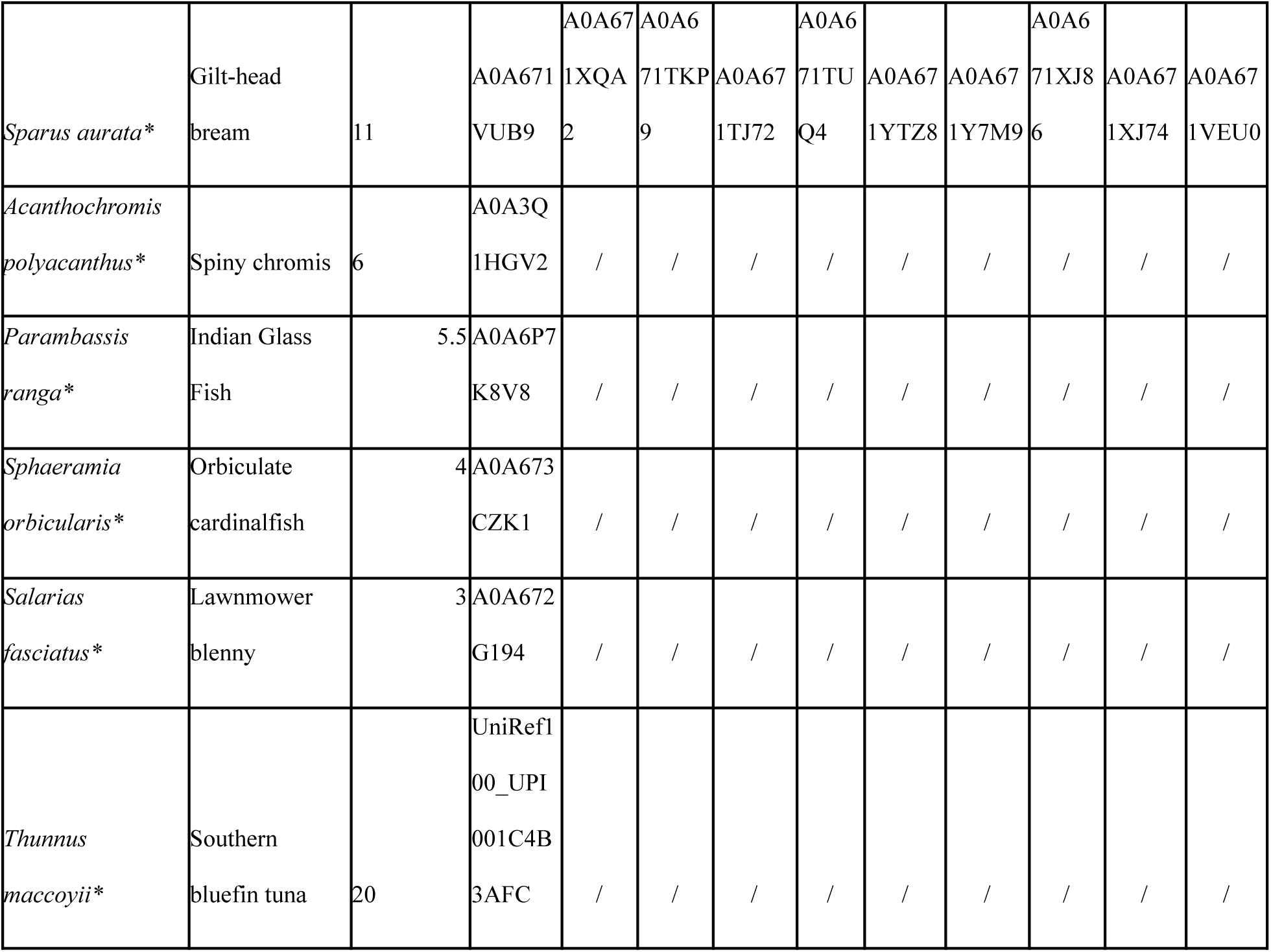
Perciformes included in the study. Organisms in order Perciformes included in the RES, SIFT, and/or PEPPI studies, their average lifespans, and UniPro or NCBI GenBank’s protein accession numbers. * indicates those organisms whose p53 sequences are used in alignment for RES study.

**S3 Table.**
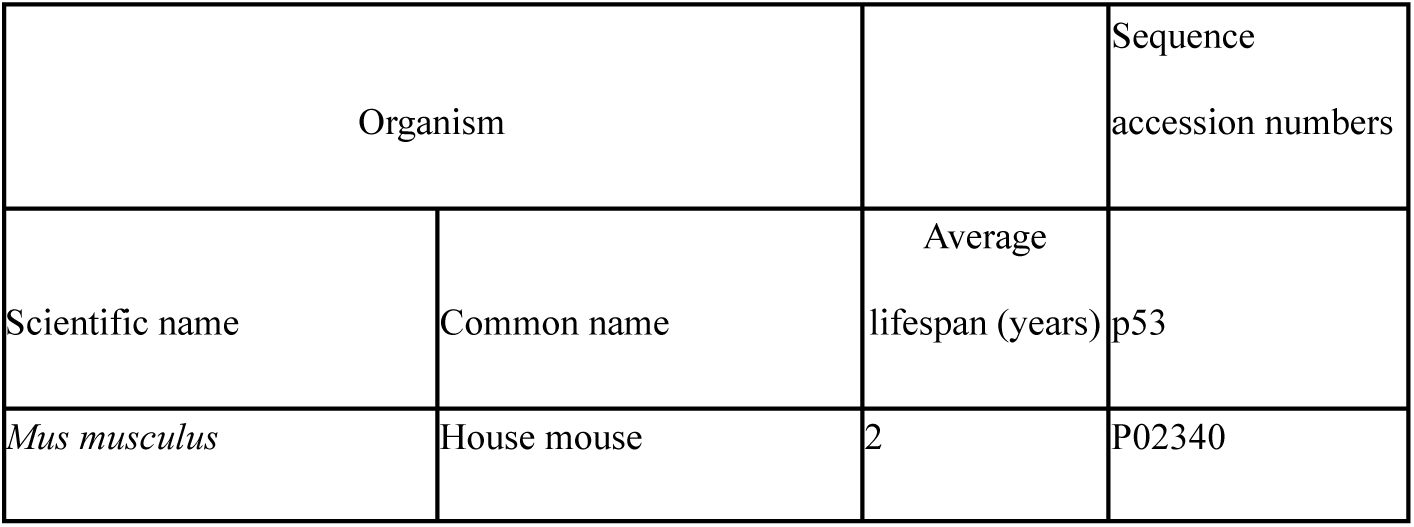
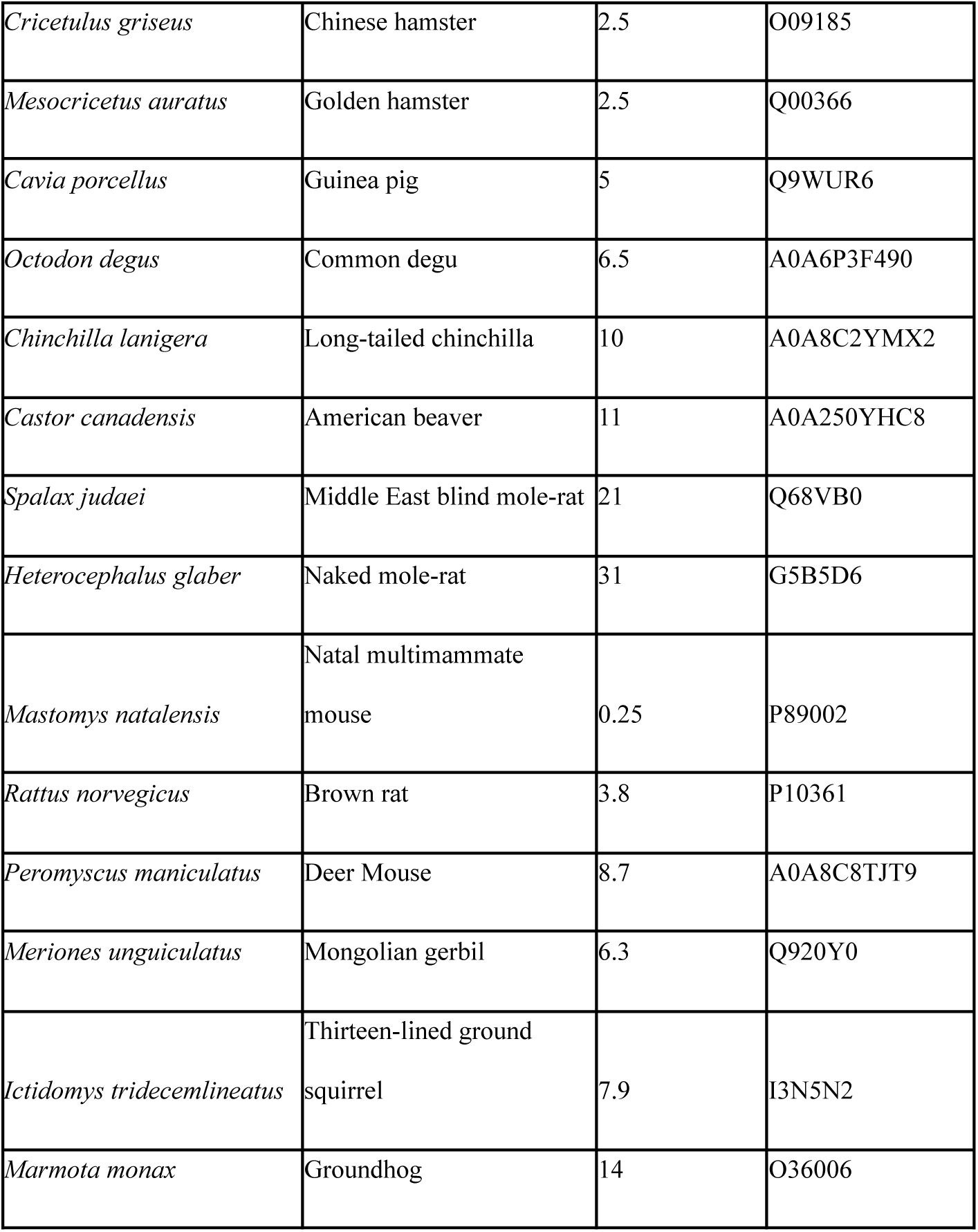
Rodents included in the study. Organisms in order Rodentia included in the RES and SIFT studies, their average lifespans, and UniPro or NCBI GenBank’s protein accession numbers.

**S4 Table.** Carnivores included in the study. Organisms in order Carnivora included in the RES and SIFT studies, their average lifespans, and UniPro or NCBI GenBank’s protein accession numbers.

| Organism |  |  | Sequence accession numbers |
| --- | --- | --- | --- |
| Scientific name | Common name | Average lifespan (years) | p53 |
| *Neovison vison* | American mink | 11.4 | A0A8C7A9R2 |
| *Vulpes vulpes* | Red fox | 12 | A0A3Q7TIN0 |
| *Canis lupus dingo* | Dingo | 14 | A0A8C0K1L6 |
| *Nyctereutes procyonoides* | Common raccoon dog | 15.6 | A0A811ZM26 |
| *Canis lupus familiaris* | Dog | 20.6 | Q29537 |
| *Neomonachus schauinslandi* | Hawaiian monk seal | 25 | A0A2Y9HEV7 |
| *Callorhinus ursinus* | Northern fur seal | 25 | A0A3Q7NG00 |
| *Panthera leo* | Lion | 28 | A0A8C8Y413 |
| *Felis catus* | Cat | 30 | P41685 |
| *Ursus maritimus* | Polar bear | 38.2 | A0A384BVC2 |
| *Zalophus californianus* | California sea lion | 30 | A0A6J2FNY3 |
| *Lynx rufus* | Bobcat | 32.3 | UPI001F1287F1 |
| *Enhydra lutris* | Sea otter | 19 | A0A2Y9L4Q2 |
| *Panthera pardus* | Leopard | 22 | A0A9V1DWE6 |
| *Suricata suricatta* | Meerkat | 12.5 | A0A673UVH2 |

**S5 Table.** Cetaceans included in the study. Organisms in order Catacean included in the RES and SIFT studies, their average lifespans, and UniPro or NCBI GenBank’s protein accession numbers.

| Organism |  |  | Sequence accession number |
| --- | --- | --- | --- |
| Scientific name | Common name | Average lifespan (years) | p53 |
| *Lipotes vexillifer* | Baiji | 24 | A0A340X8E6 |
| *Delphinapterus leucas* | Beluga whale | 40 | Q8SPZ3 |
| *Tursiops truncatus* | Common bottlenose dolphin | 45 | A0A2U4C2U9 |
| *Monodon monoceros* | Narwhal | 50 | A0A4V5P9N3 |
| *Globicephala melas* | Long-finned pilot whale | 60 | UPI00293D9E1D |
| *Eubalaena glacialis* | North Atlantic right whale | 67 | UPI002A5A2F50 |
| *Physeter macrocephalus* | Sperm whale | 68 | A0A455C1G1 |
| *Balaenoptera musculus* | Blue whale | 85 | A0A8C0DFA9 |
| *Balaenoptera physalus* | Fin whale | 114 | A0A6A1Q3Q7 |
| *Balaenoptera acutorostrata* | Common minke whale | 57 | A0A383YUA7 |
| *Lagenorhynchus albirostris* | White-beaked dolphin | 40 | XP_059988824.1 |
| *Mesoplodon densirostris* | Blainville's beaked whale | 27 | XP_059937197.1 |
| *Delphinus delphis* | Short-beaked common dolphin | 25 | XP_059854544.1 |
| *Kogia breviceps* | Pygmy sperm whale | 17 | XP_058904949.1 |
| *Phocoena phocoena* | Harbour porpoise | 13 | XP_065753055.1 |

**S6 Table.** Artiodactyles included in the study. Organisms in order Artiodactyl included in the RES and SIFT studies, their average lifespans, and UniPro or NCBI GenBank’s protein accession numbers.

| Organism |  |  | Sequence accession number |
| --- | --- | --- | --- |
| Scientific name | Common name | Average lifespan (years) | p53 |
| *Ovis aries* | Sheep | 11 | P51664 |
| *Sus scrofa* | Wild boar | 12 | Q9TUB2 |
| *Muntiacus muntjak* | Southern red muntjac | 17 | A0A5N3WLG9 |
| *Bos taurus* | Cattle | 20 | P67939 |
| *Capra hircus* | Goat | 20.8 | A0A452G0A3 |
| *Odocoileus virginianus texanus* | Texas white-tailed deer | 23 | A0A6J0VGP1 |
| *Vicugna pacos* | Alpaca | 25 | A0A6I9IU1 |
| *Bison bison bison* | Plains bison | 33.5 | A0A6P3HQA5 |
| *Camelus bactrianus* | Bactrian camel | 35 | A0A9W3H045 |
| *Hippopotamus amphibius kiboko* | Hippopotamus | 61 | XP_057572051.1 |
| *Camelus dromedarius* | Dromedary | 45 | A0A5N4D2L6 |
| *Bubalus bubalis* | Domestic water buffalo | 34.9 | F6MDM8 |
| *Bos mutus grunniens* | Wild yak | 26.3 | A0A0N7FDT7 |
| *Neophocaena asiaeorientalis asiaeorientalis* | Yangtze finless porpoise | 21.5 | A0A341BQX3 |
| *Bos indicus* | Zebu | 20 | P67938 |

RES GitHub repository can be accessed at https://github.com/Romani20/RES. It includes the RES program, installation instructions, and fasta files to score p53 sequences involved in study.

